# Low sperm to egg ratio required for successful in vitro fertilisation in a pair-spawning teleost, Senegalese sole (*Solea senegalensis*)

**DOI:** 10.1101/2020.08.26.267740

**Authors:** Sandra Ramos-Júdez, Wendy Ángela González-López, Jhons Huayanay Ostos, Noemí Cota Mamani, Carlos Marrero Alemán, José Beirão, Neil Duncan

**Author notes:** Corresponding author: Neil Duncan. These two authors made a similar contribution to the article (Sandra Ramos-Júdez, Wendy Ángela González-López).

## Abstract

Cultured Senegalese sole (*Solea senegalensis*) breeders fail to spawn fertilised eggs and this bottleneck could be solved with the implementation of large-scale *in vitro* fertilisation protocols. However, low production of poor-quality sperm has frustrated the development of *in vitro* fertilisation protocols. Cultured females were induced to ovulate with a 5 µg kg^-1^ single injection of gonadotropin releasing hormone agonist (GnRHa) and good quality eggs (82.6 ± 9.2% fertilisation) were stripped 41:57 ± 1:46 h after the injection. Sperm was collected from cultured males, diluted in modified Leibovitz and used fresh to fertilise the eggs. A non-linear regression, an exponential rise to a maximum (R = 0.93, P < 0.0001) described the number of motile spermatozoa required to fertilise a viable egg and 1617 motile spermatozoa were sufficient to fertilise 99 ± 12% (± 95% CI) of viable eggs. Similar, spermatozoa egg^-1^ ratios of 592 ± 611 motile spermatozoa egg^-1^ were used in large-scale *in vitro* fertilisations with 190,512 ± 38,471 eggs. The sperm from a single male (145 ± 50 µL or 8.0 ± 6.8 × 10^8^ spermatozoa) was used to fertilise the eggs. The mean hatching rate of the large-scale *in vitro* fertilisations was 70 ± 14 % to provide 131,540 ± 34,448 larvae per fertilisation. When unfertilised eggs were stored at room temperature the percentage of viable eggs decreased gradually and indicated the sooner eggs were fertilised after stripping the higher the viability of the eggs. The collection of sperm directly into a syringe containing modified Leibovitz significantly increased the percentage of motile spermatozoa (33.4 ± 12.2 %) compared to dilution in modified Leibovitz immediately after collection (6.6 ± 4.9 %). Senegalese sole have a pair-spawning reproductive behaviour characterised by external gamete fertilisation in close proximity with no sperm competition. The low spermatozoa egg^-1^ ratio required for maximum fertilisation was consistent with this reproductive behaviour and strategy. The provision of a large-scale *in vitro* fertilisation protocol (200 µL of sperm per 100 mL of eggs) will enable the industry to operate sustainably and implement breeding programs to improve production.

## 1. Introduction

Senegalese sole (*Solea senegalensis*) is a promising emerging aquaculture species in Europe. Sole production from land based farms in Spain, Portugal, France and Iceland has increased rapidly to 1,700 t in 2019 (APROMAR, 2019). This increase is driven by good market prices, high market demand and successful culture practices (Morais et al., 2016) that permit cost effective production despite of the need for high levels of investment in culture infrastructure.

However, the production cycle is not fully controlled and relies on the capture of wild broodstock that spawn sufficient eggs to achieve targeted aquaculture productions (Anguis and Cañavate, 2005; Martín et al., 2014). The progeny of these wild broodstock and in particular the males exhibit a reproductive behavioural dysfunction and do not participate in the courtship to fertilise eggs (Duncan et al., 2019; Fatsini et al., 2020; Guzmán et al., 2008; Martín et al., 2020). Consequentially, cultured broodstocks that were reared entirely in captivity produce unfertilised eggs (Duncan et al., 2019; Guzmán et al., 2008). Differences between wild and cultured breeders have suggested that the reproductive dysfunction has a bases in a combination of endocrine reproductive control (Guzmán et al., 2011; Riesco et al., 2019), social conditions during rearing (Fatsini et al., 2020; Martín et al., 2020), broodstock nutrition (Norambuena et al., 2013a, 2013b, 2012b, 2012c, 2012a) and olfactory capacity (Fatsini et al., 2017). However, no practical approach has been developed to overcome these reproductive dysfunctions and the low levels of fertilised egg obtained from cultured broodstocks (Fatsini et al., 2020; Guzmán et al., 2011) were insufficient to meet industry needs.

In flatfish culture, *in vitro* fertilisation methods are commonly used to obtain the fertilised eggs required for aquaculture (Mañanós et al., 2008). *In vitro* fertilisation enables aquaculturists to bypass behavioural reproductive dysfunctions of the type observed in cultured male Senegalese sole. However, the application of *in vitro* fertilisation methods for Senegalese sole has been frustrated by the small quantities of poor quality sperm produced by males (Beirão et al., 2011, 2009; Cabrita et al., 2011, 2006; González-López et al., 2020). Rasines et al. (2013, 2012), has described *in vitro* fertilisation procedures for Senegalese sole on an experimental scale. Female sole were induced to ovulate with gonadotropin releasing hormone agonist (GnRHa) and batches of 1 mL of stripped eggs were fertilised with 30 µL of cryopreserved sperm from cultured males. Similarly, Liu et al. (2008) described *in vitro* fertilisation of Senegalese sole eggs on an industrial scale. Again, eggs were obtained from GnRHa induced cultured females and all the eggs from each female were fertilised *in vitro* with the sperm from three to four cultured males. However, few details were given on the amount of sperm or eggs used and the spermatozoa (spz) to egg ratio was not detailed.

When sperm is limiting, it is of critical importance to know the spz to egg ratio (amount of spz required to fertilise each egg) to plan *in vitro* fertilisation procedures. Spermatozoa to egg ratios for *in vitro* fertilisation of fish eggs show considerable variation ranging from × 10^3^ to × 10^6^ (Beirão et al., 2019). However, it would appear that flatfish have lower spz requirements as winter flounder (*Pseudopleuronectes americanus*) required 3.4 × 10^4^ spz egg^-1^ (Butts et al., 2012) and turbot (*Scophthalmus maximus*) required just 3000 to 6000 spz egg^-1^ (Chereguini et al., 1999; Suquet et al., 1995). Many flatfish species have reproductive strategies and behaviours to spawn as a pair (Carazo et al., 2016; Gibson et al., 2014). Monogamous fish species (classified as species that spawn in a pair) were shown to have smaller testes compared to polyandrous species (group spawning of males with a female) (Baker et al., 2020). Monogamy or pair-spawning will also reduce sperm competition (sperm from two or more males compete to fertilise the eggs) and decreasing sperm competition has been related to smaller testes and lower sperm production (Parker and Pizzari, 2010; Stockley et al., 1997). Senegalese sole spawn as pairs (Carazo et al., 2016; Duncan et al., 2019) and spawning pairs show a degree fidelity during and between spawning seasons (Fatsini et al., 2020; Martín et al., 2014). Therefore, pair-spawning and low sperm competition would appear to explain the small testes size (García-López et al., 2005; González-López et al., 2020) and low sperm production reported for Senegalese sole (Beirão et al., 2011, 2009; Cabrita et al., 2011, 2006; González-López et al., 2020). In addition, a preliminary study found that Senegalese sole achieved a high percentage of fertilisation with a low spz to egg ratio (Marrero-Alemán et al., 2019) and indicated that *in vitro* fertilisation with the low numbers of sperm may be a viable solution to the industries problem to control the reproduction.

The present study, aimed to determine the spz to egg ratio required for *in vitro* fertilisation in Senegalese sole. The first aim was to determine the sperm to egg ratio on an experimental scale and then use similar ratios for commercial large-scale *in vitro* fertilisations as a proof-of-concept. Additional aims were, to determine the viability of ovulated eggs stored at room temperature and to improve sperm collection methods.

## 2. Methods

### 2.1 Experimental animals

All Senegalese sole broodstock used were cultured fish that had been hatched and reared entirely in captivity. Females used had an average weight of 1.53 ± 0.28 kg and males had a weight of 1.05 ± 0.25 kg. Fish were maintained in 10,000 L tanks in IRTA Sant Carles de la Rápita (Catalonia, Spain). Prior to experiments, fish were held in surface sea water (∼35 ppt, >5 mg.L^-1^ O2) and a controlled natural temperature cycle (9-20 °C) using recirculation systems (IRTAmar®). Tanks were covered with shade netting and photoperiod was natural with natural light. The fish were fed four days a week with either unfrozen polychaetes and mussels (0.75 % of biomass) or 5 mm pelleted Broodfeedlean broodstock diet (0.55% of biomass) (Sparos, Olhão, Portugal). During experiments conducted from April to June, fish were held in the same conditions with the exceptions that water temperature was maintained at a constant 16 ± 1 °C and fish were not feed 24 hours before any manipulation.

The fish were handled (routine husbandry and experimentation) in accordance with European regulations on animal welfare (Federation of Laboratory Animal Science Associations, FELASA, http://www.felasa.eu/). For all handling and sampling, fish were anesthetised with 60 mg L^-1^ tricaine methanesulfonate (MS-222; Sigma-Aldrich, Spain).

### 2.2 Gametes

Eggs were obtained by inducing ovulation. Ovarian biopsies were taken from females with swollen ovaries and the diameter of 20 oocytes were measured (x40 Carl Zeiss Axiostar microscope). Females were selected that had mean oocyte diameter ≥ 600 µm (Marrero-Alemán et al., 2019). The females were administered 5 µg kg^-1^ of GnRHa (Sigma code L4513, Sigma, Spain) (Agulleiro et al., 2006) between 18:00 to 19:00 h. The females were held with constant temperature (16 ± 1 °C) and total darkness until ovulation. Females were checked for ovulation every 2-3 hours starting from 40 h after the administration of GnRHa (Marrero-Alemán et al., 2019; Rasines et al., 2013, 2012) and all eggs were stripped from the ovulated females. Percentage fertilisation when spz were in excess was used to indicate egg quality.

Sperm was obtained from males by repeatedly, gently massaging the testes and applying pressure along the full length of the sperm ducts to the urogenital pore. All sperm with urine contamination was collected in a 1 mL syringe. González-López et al. (2020), demonstrated that avoiding urine contamination was almost impossible and that high numbers of motile spz were obtained stripping sperm mixed with urine. The volume collected was measured with the syringe to an accuracy of 10 µL and the sperm was transferred to a 1.5 mL Eppendorf and immediately diluted with modified Leibovitz (González-López et al., 2020) using the dilution required for the experiment (see below). The sperm motility was initially observed (x 100 Carl Zeiss Axiostar microscope) by activating 1 µL of diluted sperm with 19 µL of clean seawater. Sperm samples with low or no motility were rejected. All sperm samples were stored over ice or at 4°C (refrigerated) until analysis or used to fertilise eggs.

The spz concentration (spz mL^-1^) was measured using a Thoma cell counting chamber. A 10 µL sample of sperm was diluted 1:500 in 10% formalin and 10 µL of the dilution was pipetted into the counting chamber. After 10 minutes for spz to sediment, the chamber was observed using a microscope (x100 magnification with Olympus BH microscope), photographed (IC Capture software and GigE digital camera model: DMK 22BUC03 Monochrome, The Imaginsource, Bremen, Germany) and the number of spz counted (ImageJ software, http://imagej.nih.gov/ij/).

Sperm motility parameters were determined as described by González-López et al. (2020). Spermatozoa were activated by mixing 1 µL of diluted sperm (1:4 with modified Leibovitz) with 20 µL of seawater with 30% bovine serum albumin (BSA, Sigma, Spain). One µL of activated sperm was pipetted into an ISAS R2C10 counting chamber (Proiser *R+D, S*.*L*. Paterna, Spain) previously mounted and focused on the microscope (200x magnification Olympus BH). Video recording was initiated when spz were activated and tracks were recorded (IC Capture software and GigE digital camera) until motion ceased. Videos (AVI format) of spz tracks from 15 to 17 s (unless otherwise stated) after activation were converted into image sequences (jpeg format using Virtual Dub 1.10.4 software http://www.virtualdub.org/). The image sequences were analysed using ImageJ software with the computer-assisted sperm analysis (CASA) plugin (ImageJ http://rsb.info.nih.gov/ij/plugins/) using the settings: brightness and contrast, -10 to 15/224 to 238; threshold, 0/198 to 202; minimum sperm size (pixels), 10; maximum sperm size (pixels), 400; minimum track length (frames), 10; maximum sperm velocity between frames (pixels), 30; frame rate, 30; microns/1000 pixels, 303; Print motion, 1; the additional settings were not modified. The parameters, percentage of motile spz (% motility), Curvilinear Velocity (VCL, µm/s) and Average Path Velocity (VAP, µm/s) were recorded. All sperm samples were analysed in triplicate.

The experiments (unless otherwise stated) aimed to use gametes (eggs and sperm) as soon as possible after collection to avoid possible loses of viability due to storage. Researchers worked as two groups to strip eggs and sperm at the same time and complete *in vitro* fertilisations soon afterwards.

### 2.3 Spermatozoa to egg ratio experiment

Five different females and five different males were used during this experiment. When an ovulated female was encountered, sperm was collected and checked to find a male with ≥ 300 µL of sperm that exhibited motility. The sperm was serially diluted with modified Leibovitz (González-López et al., 2020) to achieve eight dilutions: 1:4; 1:19; 1:79; 1:319; 1:959; 1:2879; 1:5759; 1:11519. A sample of the first dilution (1:4) was used to determine the spz concentration and percentage motility. The spz concentration in the dilution 1:4 was used to calculate the spz concentration in each dilution and spz motility to calculate the concentration of motile spz. The eggs and diluted sperm were used to make three triplicate fertilisations for each serial dilution. Fertilisations were made in 100 mL beakers by pipetting in close sequence, 0.5 mL of eggs, 20 µL of diluted sperm and 5 mL clean seawater. A 1 mL pipette with a cut tip was used to pipette eggs and a 100 µL pipette with a cut tip was used to pipette diluted sperm. The eggs, sperm and seawater were gently mixed by rocking and swirling the beaker. After 3-5 minutes, the volume of seawater was topped up to 100 mL. The beakers of fertilised eggs were transferred to a 16 °C incubator. After 24 hours the eggs from each beaker were concentrated in a sieve and placed in a 10 mL Bogorov camber and ≥ 50 eggs were randomly examined using a binocular microscope (Nikon C-DSS230) to determine the number of developing eggs. In addition, the number of eggs in three 0.5 mL samples was counted for each female.

### 2.4 Proof-of-concept experiment

Seven different females and seven different males were used during this experiment. The experiment aimed to make large-scale *in vitro* fertilisations using spz egg^-1^ ratios from the previous experiment to fertilise the number of eggs (> 100,000 eggs) that would be required in a commercial fish farming scenario. When an ovulated female was encountered, sperm was collected and checked to find a male with ≥ 150 µL of motile sperm. The sperm was immediately diluted 1:4 in modified Leibovitz and a sample of 50 µL of diluted sperm was taken to determine spz concentration and percentage motility (CASA). All the eggs were stripped from the female into a clean, dry 1 L jug and the volume of eggs was measured with an accuracy of 10 mL. Three samples of 0.5 mL of eggs were taken and counted. The remaining sperm obtained from the male was added to the eggs followed by a volume of seawater that was equal to the volume of eggs. The eggs, sperm and seawater were gently swirled to mix the contents. After 2-3 minutes the jug was topped up to 1 L with seawater. The eggs were then divided into two or three parts and each part was placed in a 30 L incubator with the same conditions as the broodstock holding tanks. The number of eggs in each incubator was estimated by mixing the incubator homogenously and taking three 100 mL samples and counting the eggs in each sample. The eggs were left two days to hatch and the number of hatched larvae in each incubator was estimated as above for the eggs. The hatch rate was calculated from the number of eggs stocked and number of larvae hatched and the mean was calculated for the replica incubators used for each female - male pair.

### 2.5 Egg viability experiment

Three different females and three different males were used during this experiment. When an ovulated female was encountered, males were checked to find a male with ≥ 100 µL of motile sperm. The sperm was immediately diluted 1:4 in Leibovitz. All the eggs were stripped from the female into a clean, dry 1 L jug. The eggs were covered and stored at room temperature inside a building (out of sun light). As soon as possible after the gametes were stripped the first fertilisation was completed as previously described. The time the eggs were stripped and the time the first fertilisation was made was recorded. Further fertilisations were completed at 30 to 60 minute intervals. Fertilisations were completed in duplicate or triplicate using 0.5 mL of eggs and 20 µL of diluted sperm that ensured an excess of motile spz per egg. As described above, the beakers of fertilised eggs were transferred to a 16 °C incubator and after 24 hours the percentage of developing eggs was determined for each fertilisation.

### 2.6 Sperm collection experiment

Thirteen males were used for this experiment. Sperm from each male was collected as previously described, with the exception that the sperm was collected either into an empty clean syringe (100 µL of sperm) or into a syringe that contained modified Leibovitz (to give a 1:4 dilution, 50 µL of sperm collected into 200 µL of Leibovitz). Both collection methods were used for each male and the sequence of collection was alternated. For seven animals sperm was first collected into a clean syringe and then into a syringe that contained modified Leibovitz and for six animals the reverse, first directly into modified Leibovitz and then a clean syringe. The 100 µL of sperm collected into a clean syringe was immediately divided into 50 µL that was diluted in 200 µL of Leibovitz (1:4 dilution) and 50 µL that was kept as undiluted sperm as a control. The time of collection and dilution after collection was recorded. The sperm motility parameters were analysed (CASA) for the samples collected by these three methods: collected directly into Leibovitz, collected before dilution in Leibovitz and undiluted sperm. Sperm motility parameters were analysed 30 s after activation for 2 s. The parameters were measured at the time of collection (0 h), six and 24 h after collection.

### 2.7 Data analysis and statistics

All means are with one standard deviation unless otherwise stated. For the spermatozoa egg^-1^ ratio experiment, the percentage of viable eggs fertilised was calculated by dividing the actual fertilisation rate by the mean fertilisation rate when sperm was in excess. The number of motile sperm was calculated by multiplying the volume of diluted sperm added by the spermatozoa concentration and the percentage motility. To examine the effect that gamete quality amongst the five pairs of fish had on fertilization, a linear regression was applied to percentage of motile sperm (sperm quality) against number of sperm (motile and immotile) required per viable egg and to percentage of viable eggs (egg quality) against number of motile sperm required per egg. To describe the variation of percentage of viable eggs fertilised in relation to the number of motile spz per egg, a non-linear regression based on an equation for an exponential rise to a maximum with double, five parameters was applied to the data. For the egg viability experiment, the percentage of viable eggs fertilised was calculated as above. To describe the variation of percentage of viable eggs fertilised in relation to time the eggs were stored at room temperature a non-linear regression based on an equation for a four parameter logistic curve was applied to the data.

The data set for percentage motility from the sperm collection experiment was not normally distributed with a high percentage of zeros and skewed positively to a few higher values. The data set could not be transformed to normality. The data was analysed twice. In one analysis, the data from the time point zero was ranked and analysed with a two-way ANOVA with the independent variables order of collection (1^st^ or 2^nd^) and collection treatment (collected directly into Leibovitz, diluted in Leibovitz and undiluted sperm). In a second analysis the data set was scored into samples with or without motility and a Chi squared analysis was make to compare expected proportions of samples with motility with actual proportions with motility between sperm collection treatments (collected directly into Leibovitz, diluted in Leibovitz and undiluted sperm) and time of storage (0 h, 6 h and 24 h). The Marascuillo procedure (Prins, 2012) was used to make a multiple comparison between proportions of individual treatments and time points. All the samples with motility were then separated (i.e. all the zeros were excluded) into a smaller data set that was transformed to normality with the Logit transformation. The transformed data was compared with a one way ANOVA followed by Holm-Sidak pairwise multiple comparison to compare mean motility for each treatment at each time point. A P<0.05 was used to indicate significant differences. All statistical comparisons and regressions were made using Sigma Plot 12 (Systat Software, Inc., San Jose, CA 95110. USA) except for the Marascuillo procedure that was completed with an Excel (Microsoft) worksheet written by the authors.

## 3.0 Results

A total of 46 males were checked to obtain 28 (60.9 %) males with the required sperm quantity and quality for the experiments. All males were only used once. A total of 20 females were selected by ovarian swelling and oocyte diameter determined. Five females were rejected as the ovaries contained ovulated ova and two as the ovaries had solid cysts. A total of 13 females had oocytes ≥ 600 µm and were induced with GnRHa. Five females were not used as four females did not ovulate and one female had low quality eggs (0% fertilisation). Eight (62%) females ovulated good quality eggs that were used in the different experiments. The eggs from some females were used for more than one experiment. The mean latency time from injection with GnRHa to the ovulation was 41:57 ± 1:46 h and mean fertilisation was 82.6 ± 9.2%. The mean fecundity was 130,789 ± 36,723 eggs fish^-1^ or 87,174 ± 24,378 egg kg^-1^ of female body weight. The maximum time difference between stripping eggs and sperm was 30 minutes, therefore, either sperm or eggs were stored for 30 min or less before fertilisation experiments were initiated.

### 3.1 Spermatozoa to egg ratio experiment

The percentage of viable eggs fertilised in relation to number of motile spz per egg showed a rapid increase from zero that was represented by a non-linear regression based on an equation for an exponential rise to a maximum with double, five parameters (R = 0.93, P < 0.0001) (Fig. 1). The non-linear regression described that only 326 motile spz per egg fertilised 79 ± 15% (± 95% CI - confidence interval) of viable eggs, that 649 motile sperm fertilised 90 ± 13% (± 95% CI) of viable eggs and 1617 motile spz fertilised 99 ± 12% (± 95% CI) of viable eggs.

**Figure 1.**
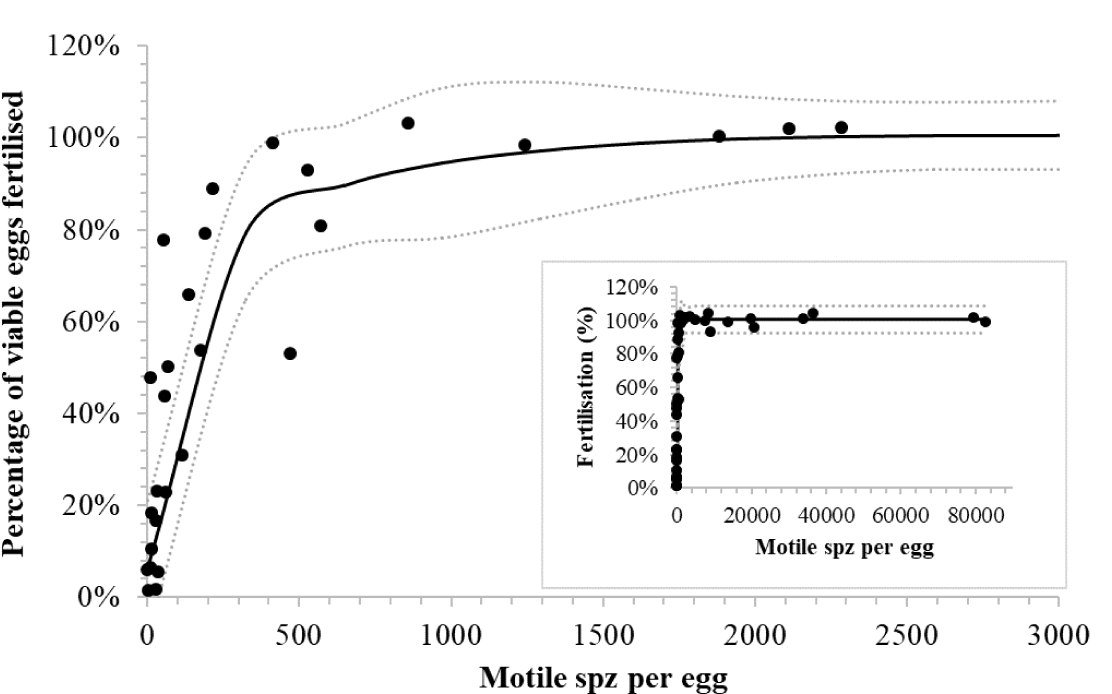
The percentage of viable eggs fertilised in relation to the number of motile spermatozoa (spz) per viable egg for Senegalese sole (*Solea senegalensis*). The insert figure shows the entire data set up to over 80,000 spz per egg and the large figure shows a close up of the data up to 3,000 motile spz per egg. The continuous line shows a non-linear regression based on an equation for an exponential rise to a maximum with double, five parameters (R = 0.93, P < 0.0001) that represents the variation in percentage of viable eggs fertilised in relation to number of motile sperm per egg. The dotted lines indicate 95% confidence intervals for the non-linear regression.

Amongst the five pairs, the percentage motility of the spz was correlated to the number of spz (motile and non-motile) required to fertilise a viable egg (R^2^ = 0.83, P = 0.021). However, there was no correlation between percentage of viable eggs and number of motile sperm required to fertilise each egg (R^2^ = 0.03, P = 0.37). Caution is required in the interpretation as n was low (n = 5) and the statistical power (at α = 0.05) of the tests was 0.66 and 0.13, respectively for percentage motility and viable eggs.

### 3.2 Proof-of-concept experiment

The proof-of-concept large-scale *in vitro* fertilisations (n = 7) gave a mean percentage hatch of 70 ± 14 % to produce a mean of 131,540 ± 34,448 larvae per fertilisation (Table 1). The sperm from a single selected cultured male with a volume of 145 ± 50 µL and total spz count of 8 ± 6.8 × 10^8^ was sufficient to fertilise large numbers of eggs (190,512 ± 38,471) and produce large numbers of larvae (131,540 ± 34,448). The mean number of spz per egg used for the commercial fertilisations was 2,981 ± 2,932 spz egg^-1^ or 592 ± 611 motile spz egg^-1^.

**Table 1.** Data from seven commercial scale *in vitro* fertilisations made for Senegalese sole (*Solea senegalensis*). Latency time from application of GnRHa (5 µg kg ^-1^) to stripping of ovulated eggs, volume of eggs used, total number of eggs used, volume of sperm used (sperm from single male before dilution), total number of spermatozoa (spz) added, number of spz per egg added, percentage motility of spz, number of motile spz per egg, mean percentage hatch in incubators and total number of larvae produced.

| Female | Latency time | Vol. eggs (mL) | Total n° eggs | Vol. sperm (µL) | Total n° Spz. x10<sup>8</sup> | N° spz. per egg | Percentage spz motility | Total n° motile spz. per egg | Percentage hatch ± SD | Total n° larvae |
| --- | --- | --- | --- | --- | --- | --- | --- | --- | --- | --- |
| 1 | 41:20 | 120 | 161,280 | 140 | 14.0 | 8,681 | 10.4 | 905 | 82 ± 16 | 132,415 |
| 2 | 43:15 | 150 | 252,450 | 140 | 3.3 | 129 | 21.9 | 28 | 79 ± 14 | 198,530 |
| 3 | 40:45 | 75 | 144,250 | 100 | 6.9 | 4,802 | 38.1 | 1830 | 78 ± 24 | 112,600 |
| 4 | 42:00 | 110 | 181,940 | 120 | 1.5 | 824 | 30.0 | 247 | 58 ± 10 | 104,625 |
| 5 | 39:30 | 100 | 198,467 | 160 | 3.5 | 1,747 | 22.2 | 388 | 77 ± 7 | 153,245 |
| 6 | 43:40 | 100 | 167,500 | 120 | 4.9 | 2,925 | 8.3 | 242 | 71 ± 11 | 117,500 |
| 7 | 44:40 | 150 | 227,700 | 120 | 4.0 | 1,757 | 28.8 | 507 | 44 ± 9 | 101,865 |
| **Mean** | **41:57** | **115** | **190,512** | **145** | **8.0** | **2,981** | **22.8** | **592** | **70** | **131,540** |
| ± SD | 01:46 | 27.5 | 38,471 | 50 | 6.8 | 2,933 | 10.7 | 611 | 14 | 34,448 |

### 3.3 Egg viability experiment

The percentage of viable eggs fertilised decreased gradually from after being stripped (Fig. 2). The non-linear regression based on an equation for a four parameter logistic curve (R = 0.80, P = 0.008) represented the variation in percentage of viable eggs fertilised in relation to the time eggs from the three females were stored at room temperature. The non-linear regression indicated that after 30 min, the percentage fertilisation had decreased from 100% to 81 ± 26% (± 95% CI), after an hour to 57 ± 20% (± 95% CI) and after two hours 32 ± 19% (± 95% CI) fertilisation of viable eggs.

**Figure 2.**
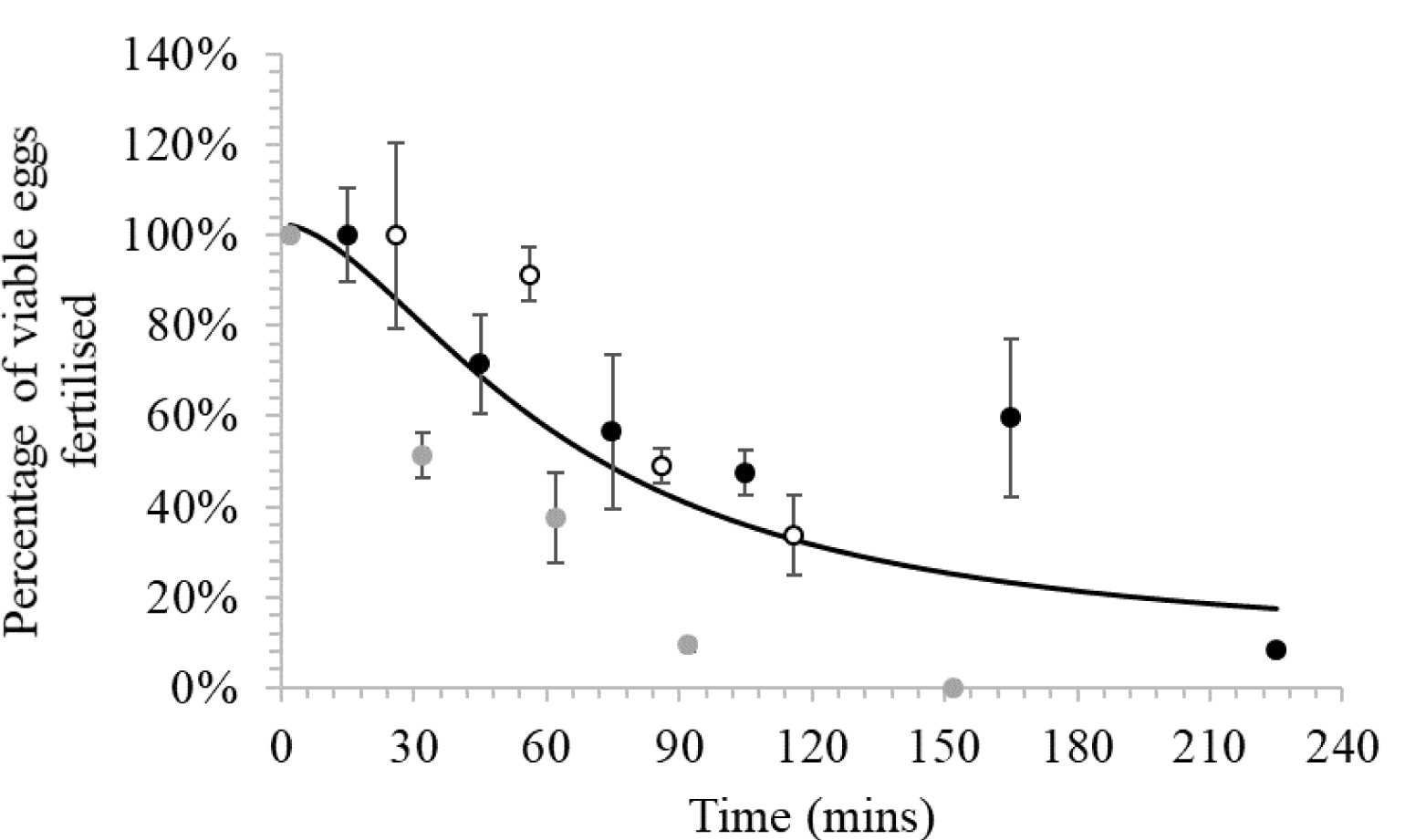
The percentage of viable eggs fertilised in relation to time eggs were stored at room temperature for three Senegalese sole (*Solea senegalensis*) females. Different dots represent different females. The line shows a non-linear regression based on an equation for a four parameter logistic curve (R = 0.08, P = 0.008) that represents the variation in percentage of viable eggs fertilised in relation to time the eggs from the three females were stored.

### 3.4 Sperm extraction experiment

The order of collection of the sperm appeared to have a significant effect and the first part of the sperm collected had significantly (P = 0.04) higher motility than the second part (Fig. 3). However, caution is required in the interpretation as there was considerable variation and the statistical power of the test was low (at α = 0.05, power = 0.44). The percentage of samples that had motility decreased significantly (P < 0.001) with time (Fig. 4). At the time of collection (t = 0) there were no differences in the percentage of samples with motility, but after six and 24 h of storage at 4°C the samples collected directly into modified Leibovitz had significantly (P < 0.05) more samples with motility.

**Figure 3.**
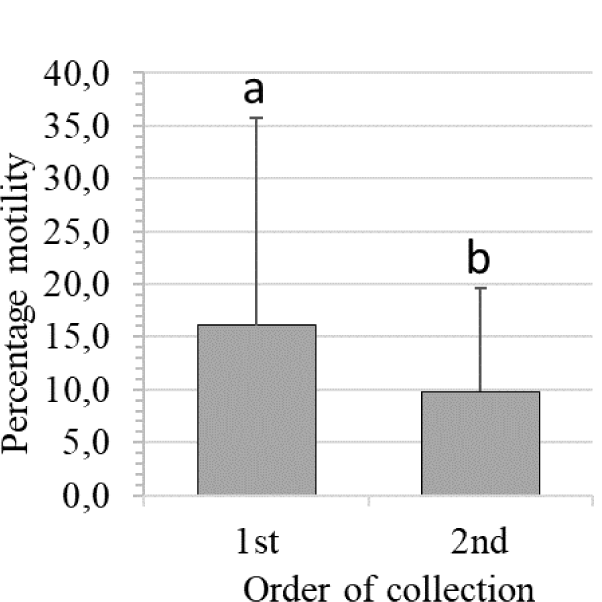
Mean percentage motility (± 1 standard deviation) of sperm samples collected first or second from each Senegalese sole (*Solea senegalensis*) male. Different letters indicate a significant difference (P = 0.04, α = 0.05, power = 0.44).

**Figure 4.**
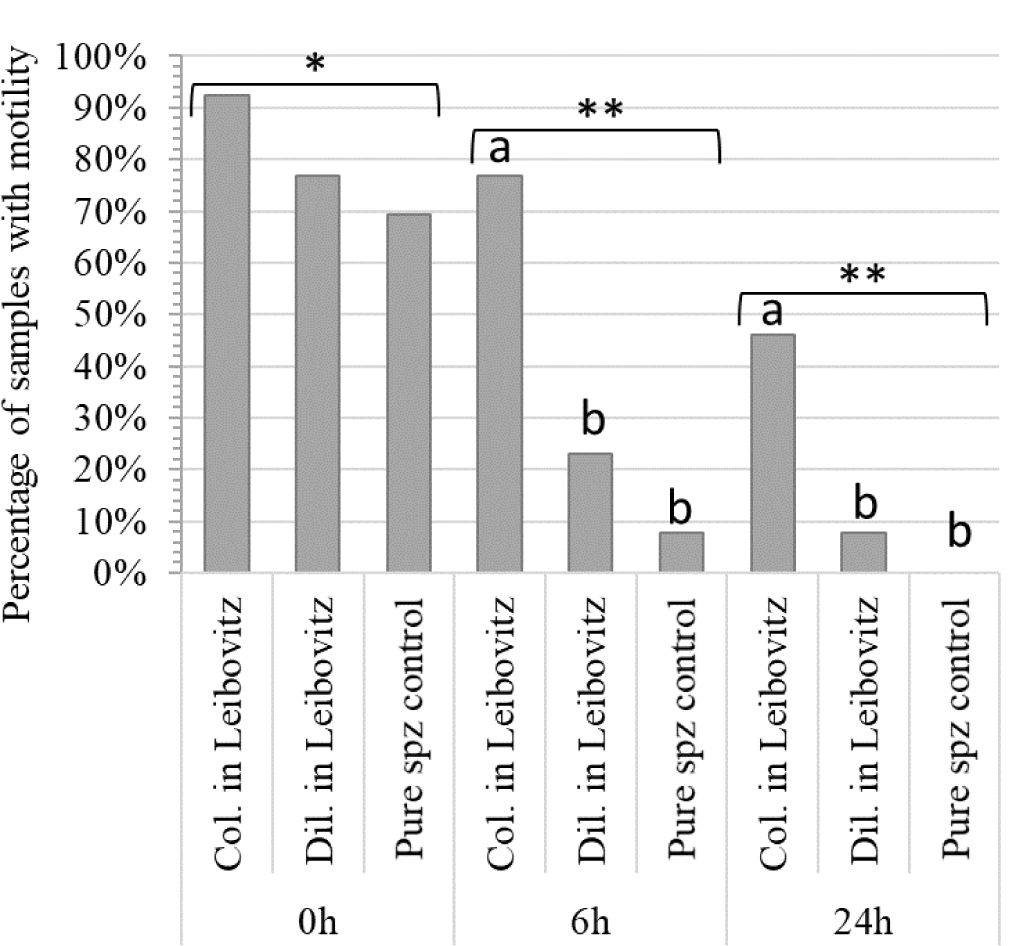
The percentage of sperm samples from Senegalese sole (*Solea senegalensis*) (n = 13) that had sperm motility for different time points 0 h after collection and after six (6 h) and 24 hours (24 h) of storage at 4°C. Three sperm collection methods were tested, collected directly into modified Leibovitz (Col. in Leibovitz), diluted in modified Leibovitz (Dil. in Leibovitz) after collection and undiluted sperm (control). Different letters indicate a significant difference (P < 0.05) within a time point and different number of asterisk indicate significant difference (P < 0.05) between time points.

The percentage motility was significantly higher (P < 0.05) in samples collected directly into modified Leibovitz at t = 0 compared to other time points (t = 6 and 24) and other collection methods, diluted in modified Leibovitz and undiluted (control), at all time points (t = 0, 6 and 24) (Fig. 5). The motility of samples collected directly into Leibovitz and stored for six hours was significantly (P < 0.05) higher than undiluted sperm samples at collection (t = 0). The time difference in mixing sperm with Leibovitz was 4 ± 2 minutes, between samples collected directly into Leibovitz and the sperm diluted in Leibovitz after collection.

**Figure 5.**
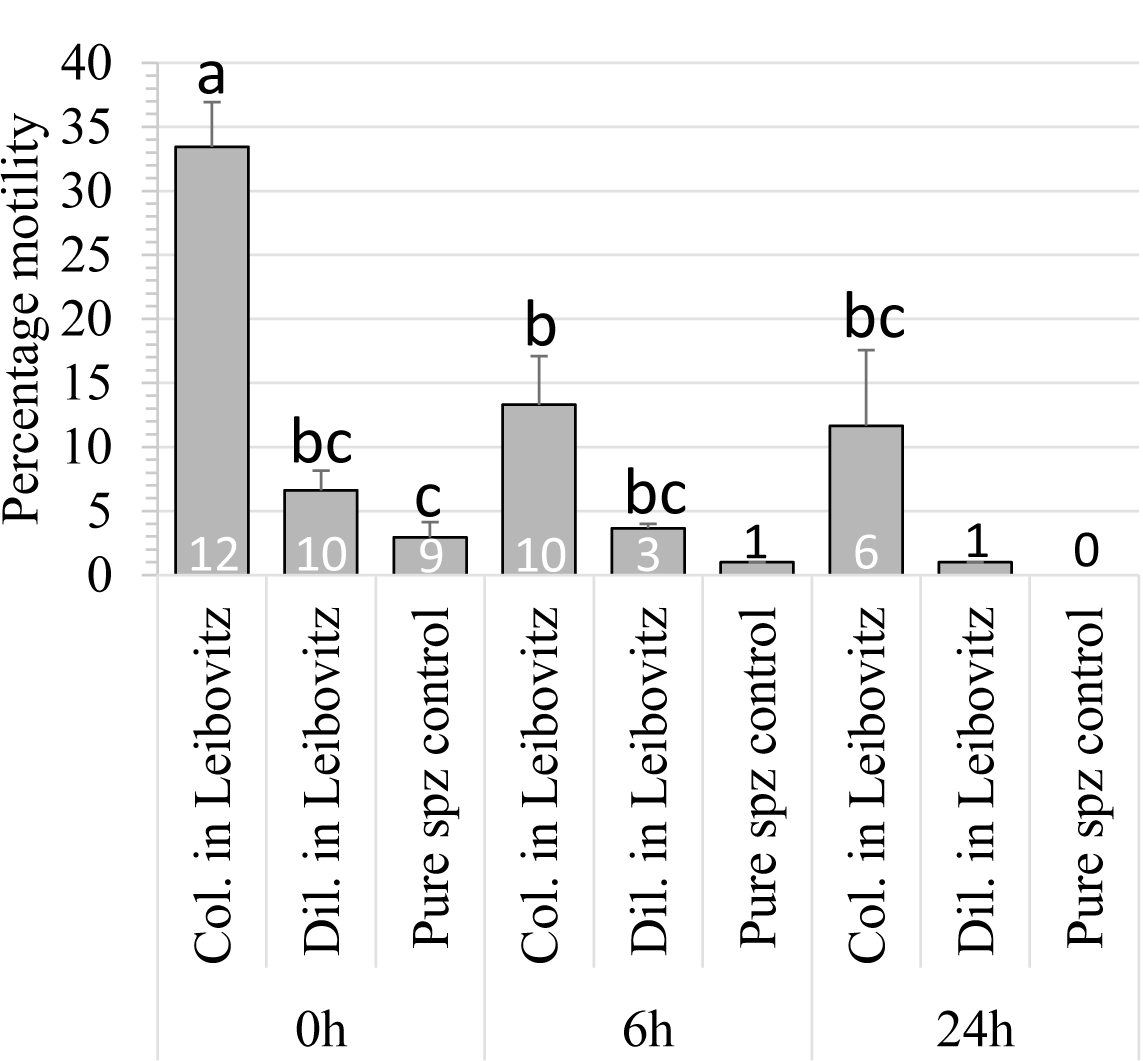
Mean percentage motility (± 1 standard error of the mean) of sperm samples collected from Senegalese sole (Solea senegalensis) using three collection methods, collected directly into modified Leibovitz (Col. in Leibovitz), diluted in modified Leibovitz (Dil. in Leibovitz) after collection and undiluted sperm (control). The motility of samples was tested at different time points 0 h after collection and after six (6 h) and 24 hours (24 h) of storage at 4°C. Different letters indicate a significant difference (P < 0.05) amongst collection methods and time points. The “n” of each mean is at the bottom of each column. Means with less than n = 3 were not included in the statistical test.

## 4.0 Discussion

### 4.1 Low spermatozoa to egg ratio in Senegalese sole

The present study demonstrated that low numbers of spermatozoa egg^-1^ ensured high levels of fertilisation in Senegalese sole. An exponential rise to a maximum (R = 0.93, P < 0.0001) described the number of motile spz required to fertilise a viable egg, the relationship rose steeply from 0 to 90% fertilisation and from approximately 500 motile spz plateaued at 90 to 100 % fertilisation. A total of 1617 motile spz per egg were sufficient to fertilise 99 ± 12% (± 95% CI) of viable eggs. A low mean spz egg^-1^ ratio also gave high rates of hatching in the proof-of-concept trials with large-scale *in vitro* fertilisations. A mean of 190,512 ± 38,471 eggs from a single female were fertilised with a mean volume of sperm of 145 ± 50 µL from a single male. This volume of sperm provided a mean of 592 ± 611 motile spz per viable egg, which was sufficient to achieve 70 ± 14 % hatch and produce 131,540 ± 34,448 larvae per *in vitro* fertilisation. The spz egg^-1^ ratios for Senegalese sole were at the lower end of ratios required for fish. Studies have shown that different fish species require a wide range of spz egg^-1^ ratios to achieve high rates of fertilisation (Beirão et al., 2019). These ratios ranged from 3000 spz egg^-1^ for turbot (Chereguini et al., 1999) to 1 × 10^5^ spz egg^-1^ for Atlantic cod (*Gadus morhua*) (Butts et al., 2009). Large-scale *in vitro* fertilisations protocols used in a hatchery generally refer to the volume of sperm needed to fertilise eggs, for example in rainbow trout 1 mL of sperm was recommended for 10,000 eggs (Bromage, 1992) and for Atlantic halibut (*Hippoglossus hippoglossus*) 1 mL of sperm for 1 L of eggs (Brown, 2010). For Senegalese sole, the present study has shown that a conservative estimation for hatcheries based on the experiments and the 95% standard deviations, would be 200 µL of sperm (≈ 2.5 × 10^8^ motile spz) to fertilise 100 mL of eggs (≈ 150,000 eggs).

### 4.2 Improved *in vitro* protocol

An aspect that complicates *in vitro* fertilisation protocols is ensuring good gamete quality for fertilisation. Generally, the sperm is obtained first and stored until eggs are obtained (Mylonas et al., 2017). Considering that Senegalese sole sperm has poor quality (González-López et al., 2020) and eggs have a short period of viability (Rasines et al., 2013, 2012), the storage of eggs and collection methods of sperm were examined in the present study to improve the *in vitro* fertilisation protocol. In addition, the present study ensured sperm or eggs were not stored for longer than 30 minutes before fertilisation to limit the effect of gamete quality deterioration. This proved to be necessary as the decline in quality of stripped Senegalese sole eggs stored at room temperature was gradual and continuous. Egg quality appeared to decline during the first 30 min of storage with no plateau period of good egg quality, which indicated the sooner eggs were fertilised after stripping the higher the viability of the eggs. This decline in egg quality after ovulation and stripping has been described in a wide range of species as the overripening process, where eggs age and in association with morphological and biochemical changes lose viability and fertilisation rates decline (Mañanós et al., 2008; Ramos-Júdez et al., 2019; Samarin et al., 2011). A rapid decline in egg quality has been observed in other species, but the decline was initiated after a period of good egg quality of 1 h in curimata (*Prochilodus marggravii*) (Rizzo et al., 2003) and 50 min in meagre (*Argyrosomus regius*) (Ramos-Júdez et al., 2019). There is considerable variation across species and some species have very different egg storage capacities, for example eggs from a Cyprinidae species kutum (*Rutilus frisii*) (Samarin et al., 2011) maintained good egg quality during eight hours of storage and salmonid eggs can be stored successfully for 4-5 days (Bromage, 1992).

To improve sperm quality different methods of sperm collection were compared. The collection of sperm directly into modified Leibovitz significantly increased motility at the time of collection and the storage capacity in terms of motility and number of samples with motility. Senegalese sole sperm is difficult or impossible to collect without urine contamination (González-López et al., 2020). The urine has negative effects on the sperm quality, which as in other species appeared to change osmolality and pH, which prematurely activated spz (Cejko et al., 2010; González-López et al., 2020; Linhart et al., 2003; Perchec Poupard et al., 1998). The collection into modified Leibovitz mitigated the negative effect of urine contamination to improve the sperm quality (González-López et al., 2020). Other studies on species where urine contamination was difficult to avoid have focused on improved collection methods that reduce urine contamination with the use of catheters that were inserted into the sperm duct (Babiak et al., 2006; Sarosiek et al., 2016) or used a collecting pipette in combination with vacuum aspiration (Gallego et al., 2013a). The very small volumes and small diameter of the urogenital pore in Senegalese sole make these kinds of approaches difficult or impossible. Small volumes of contaminated sperm obtained from some species of birds were collected directly into syringes with extender to maintain sperm quality (Personal communication, Dr. Ignacio Giménez Nebot, Rara Avis Biotec S.L., Valencia, Spain). Therefore, in the present study, this approach was taken to reduce the time that spz were in contact with urine contamination before dilution in the Leibovitz extender. A significant improvement in sperm quality was obtained by collecting the sperm directly into a syringe containing modified Leibovitz, which reduced by 4 ± 2 minutes the time before sperm was diluted with Leibovitz.

### 4.3 Sustainable Senegalese sole aquaculture

The success of massive *in vitro* fertilisations, combined with indications of egg viability during storage and improved methods for collecting and managing sperm, provided a protocol that can be used on an industrial scale to obtain eggs from cultured breeders for hatchery production. At present, the Senegalese sole aquaculture industry relies on obtaining eggs from wild adult breeders captured in the commercial fishery and there is no sustainable fishery for Senegalese sole (https://fisheries.msc.org). Therefore, obtaining viable eggs from cultured breeders has been a bottleneck that makes the industry unsustainable and unable to implement breeding programs. Breeding programs are an essential part of an aquaculture business plan that enable companies to improve growth and product quality. However, ideally, reproduction must be controlled to enable the selection and production of viable gametes from any animal that has the desired production traits. In the present study, 61% of the males checked for sperm had the required quantity and quality needed for the experiments and *in vitro* fertilisations. This availability of males, has already be improved as therapies with recombinant gonadotropins exist that both increase sperm production and quality (Chauvigné et al., 2018, 2017). These recombinant gonadotropin therapies, significantly increased sperm production by up to seven times and significantly increased the sperm quality parameters, percentage motility, progressivity and velocity of spz.

In the present study, females were initially selected by ovarian swelling and it is unclear what percentage of females would be available for selection over a reproductive season. Of the females that were GnRHa-induced, 62% ovulated good quality eggs. Therefore, studies are needed to identify the number of females available for GnRHa induction and to improve the success rate of GnRHa inductions. The present study used a GnRHa dose of 5 µg kg^-1^ to induce the ovulation of eggs compared to 25 µg kg^-1^ used by Rasines et al. (2013, 2012). These studies had similar holding conditions and temperature (16 °C) and obtained very similar timing of ovulation with means close to 42 h (range of 39 to 44 h). Egg quality appeared to be higher in the present study, but differences in methods and particular sperm storage and usage make comparisons inappropriate. Different doses of GnRHa have been compared to induce spontaneous liberations of eggs in Senegalese sole. Agulleiro et al. (2006), tested the injection of GnRHa doses of one, five and 30 µg kg^-1^. The dose of 5 µg kg^-1^ produced the most eggs and the dose of 30 µg kg^-1^ produced no liberations of eggs. Guzman et al. (2009) compared injections of 5 and 25 µg kg^-1^ of GnRHa, but found no differences in number of eggs released between GnRHa injected and untreated control fish. However, the number of oocytes advancing to hydration in the females treated with 5 µg kg^-1^ appeared to be higher than in females treated with 25 µg kg^-1^ and controls. The present study, combined with studies on GnRHa induced spontaneous liberations of eggs, would indicate that the lower dose of 5 µg kg^-1^ of GnRHa provided similar ovulation timing and egg quality as 25 µg kg^-1^ and perhaps suggest that better results may be obtained with lower doses.

### 4.4 Gamete quality and fertilisation

Logically, the spz egg^-1^ ratio required is related to the success of individual sperm to fertilise an egg. The success of spz will depend on factors of the fertilisation environment and gamete quality / characteristics that hinder or aid the spz to reach the micropyle of the egg. The environment used for *in vitro* fertilisation has been shown to affect the spz egg^-1^ ratio. For example, the volume or space provided for fertilisation affected the spz egg^-1^ ratio, as larger volumes increased the space to be travelled to fertilise the egg and increased the number of spz required (Bombardelli et al., 2013; Chereguini et al., 1999; Sanches et al., 2016). Therefore, variation in the fertilisation environment complicates the comparison of different studies within and amongst species. However, as in the present study the fertilisation environment can be standardised between tests and replicated to ensure results can be compared.

Gamete characteristics and / or quality vary amongst species and individuals. Variations in gamete quality have been shown to affect the spz egg^-1^ ratio. Obviously, percentage sperm motility affects the ratio (Gallego et al., 2013b; Moccia and Munkittrick, 1987), but velocity has also been shown to affect the ratio in walleye (*Sander vitreus*) (Casselman et al., 2006) and pufferfish (*Takifugu niphobles*) (Gallego et al., 2013b). Sperm with higher percentage motility (Gallego et al., 2013b; Moccia and Munkittrick, 1987) and spermatozoa with higher mean swimming speeds had lower spz egg^-1^ ratios to fertilise a high percentage of eggs (Casselman et al., 2006; Gallego et al., 2013b). In the present study, sperm percentage motility was related positively with spz egg^-1^ ratio (R^2^ = 0.83, P = 0.021). Senegalese sole sperm has variable and generally poor quality with low percentage motility (González-López et al., 2020). In the present study as in other studies (Ramos-Júdez et al., 2019), the effect that percentage motility has on the spz egg^-1^ ratio was removed by examining the relationship between the number of motile sperm and eggs fertilised. Many studies simply express total number of sperm (including immotile sperm) in the spz egg^-1^ ratio, however, this is inaccurate and should be stated with the percentage motility of the sperm used. Although the practice of using total spz is accepted in the literature it should only be applied to species that have little variation in motility and preferably high levels (close to 100%) of motility amongst individuals.

In the present study, the quality of eggs did not appear to be related to the spz egg^-1^ ratio. However, the low variation in egg quality (82.6 ± 9.2%) and n (n=5) may have reduced the possibility to determine a relationship. Previous studies are contradictory indicating that eggs with higher quality required more (Ramos-Júdez et al., 2019) and less (Bombardelli et al., 2013) spz than low quality eggs. It would appear probable that both the quality and the characteristics of the unfertilised egg are implicated in the fertilisation success. Good quality eggs would have more eggs to be fertilised and low quality eggs would have an environment with more space to encounter viable eggs amongst the inviable eggs. In addition, fish eggs of some species have been shown to have no mechanisms to attract spz whilst other species eggs have chemical and physical properties that guide the spz to the micropyle (Yanagimachi et al., 2017). The ability to attract spz to the eggs would in theory reduce the spz egg^-1^ ratio. These observations would suggest number of motile spz per viable egg should be used to ensure egg viability does not affect the spz egg^-1^ ratio. However, further work is required to determine the effect of egg quality on the spz egg^-1^ ratio.

### 4.5 Why do Senegalese sole have a low spz egg^-1^ ratio?

Senegalese sole spawn as a female and male pair with no involvement of other individuals (Carazo et al., 2016) and spawning pairs show a degree fidelity during and between spawning seasons (Fatsini et al., 2020; Martín et al., 2014). Therefore, Senegalese sole fertilisation does not involve sperm competition as all the sperm originates from a single male and does not compete with sperm from other males. In addition, the two sexes swim in synchrony with the genital pores held close together (Carazo et al., 2016). The male urogenital duct is slightly raised and the female oviduct forms a kind of well when eggs are being stripped (personal observations), which together with the closeness of the fish during gamete liberation (Carazo et al., 2016) suggest that the male and female place the spz next to the eggs in very close proximity. Other studies have demonstrated that these behavioural and reproductive strategies are related to low spz egg^-1^ ratios or low sperm production. Reducing the space of the fertilisation environment has been shown to reduce the spz egg ratios^-1^ required (Bombardelli et al., 2013). Different species have very varied strategies and behaviours that will alter the space of the fertilisation environment, which can range from mass spawning in aggregations in open water (Domeier and Colin, 1997; Ibarra-Zatarain and Duncan, 2015) to spawning between two fish in an enclosed space or in very close proximity (Carazo et al., 2016; Tatarenkov et al., 2006). The spawning behaviour and number of individuals involved will influence the degree of sperm competition that gametes must negotiate to achieve fertilisation. Sperm competition has been shown to influence fertilisation success and the number of spz that a species produces (Parker and Pizzari, 2010; Stockley et al., 1997). Monogamy, and the absence of sperm competition was demonstrated to reduce testes size across different taxa and monogamous fish species defined as spawning in a pair had significantly smaller testes compared to polyandrous species (group spawning of males with a female) (Baker et al., 2020). Therefore, as has been suggested for fresh water fish (Kholodnyy et al., 2020), reproductive strategies and behaviour appear to be linked to gamete requirements to achieve high rates of fertilisation. Consequentially, the reproductive strategy and behaviour of Senegalese sole support the described low spz production (García-López et al., 2005; González-López et al., 2020) as well as the low spz egg^-1^ ratio for fertilisation (present study). It can be hypothesised that spz egg^-1^ ratio is related to reproductive strategies and sperm production, however, more work is required across a wide range of species to determine the existence of a relationship.

### 4.6 Conclusion

In conclusion, Senegalese sole require a low spz egg^-1^ ratio to achieve high percentages of fertilisation both on an experimental scale and in proof-of-concept large-scale *in vitro* fertilisations. The low spz egg^-1^ ratio required to fertilise all viable eggs was consistent with the reproductive behaviour and strategies of the species. The protocol (200 µL of sperm per 100 mL of eggs) described in the present study will enable the Senegalese sole aquaculture industry to operate sustainably and establish breeding programs.

## 5. Acknowledgements

The authors would like to thank Josep Lluis Celades and the IRTA staff for technical assistance. Special thanks are also given for the participation and great enthusiasm to complete work to a high standard shown by Mario Villalta Vega and Alex Rullo Reverté, two work experience students from the IES Alfacs Escola d’Aqüicultura (http://aquicultura.insalfacs.cat/). This study has been supported with funding from the Spanish National Institute for Agronomic Research (Instituto Nacional de Investigación y Tecnología Agraria y Alimentación - INIA)-European Fund for Economic and Regional Development (FEDER) (RTA2014-0048) coordinated by Neil Duncan. The study was also supported and driven by discussions in the project 038433_REARLING, funded by Portugal and the European Union through FEDER/ERDF, NORTE 2020, in the framework of Portugal 2020, coordinated by Isidro Blanquet (Sea8 Group, Povoa de Varzim, Portugal) and in collaboration with Joan Cerdá (IRTA) and Ignacio Giménez, (Rara Avis Biotec S.L., Valencia, Spain). In particular special thanks are given to Ignacio Giménez who refused an authorship after selflessly giving his advice to help improve sperm collection and storage. Participation by Wendy González-López was funded by a PhD grant from the National Board of Science and Technology (CONACYT, Mexico) and Sandra Ramos-Júdez by a PhD grant from AGAUR (Government of Catalonia) co-financed by the European Social Fund.

## References

Agulleiro, M.J., Anguis, V., Cañavate, J.P., Martínez-Rodríguez, G., Mylonas, C.C., Cerdà, J., 2006. Induction of spawning of captive-reared Senegal sole (Solea senegalensis) using different administration methods for gonadotropin-releasing hormone agonist. Aquaculture 257, 511–524. https://doi.org/10.1016/j.aquaculture.2006.02.001

Anguis, V., Cañavate, J.P., 2005. Spawning of captive Senegal sole (Solea senegalensis) under a naturally fluctuating temperature regime. Aquaculture 243, 133–145. https://doi.org/10.1016/j.aquaculture.2004.09.026

APROMAR, 2019. La acuicultura en España 2019, Asociación Empresarial de Acuicultura de España (APROMAR). APROMAR. http://www.apromar.es/content/informes-anuales Accessed 17/08/2020

Babiak, I., Ottesen, O., Rudolfsen, G., Johnsen, S., 2006. Chilled storage of semen from Atlantic halibut, Hippoglossus hippoglossus L. Theriogenology 66, 2025–2035. https://doi.org/10.1016/j.theriogenology.2006.06.003

Baker, J., Humphries, S., Ferguson-Gow, H., Meade, A., Venditti, C., 2020. Rapid decreases in relative testes mass among monogamous birds but not in other vertebrates. Ecol. Lett. 23, 283–292. https://doi.org/10.1111/ele.13431

Beirão, J., Boulais, M., Gallego, V., O’Brien, J.K., Peixoto, S., Robeck, T.R., Cabrita, E., 2019. Sperm handling in aquatic animals for artificial reproduction. Theriogenology 133, 161–178. https://doi.org/10.1016/j.theriogenology.2019.05.004

Beirão, J., Soares, F., Herráez, M.P., Dinis, M.T., Cabrita, E., 2011. Changes in Solea senegalensis sperm quality throughout the year. Anim. Reprod. Sci. 126, 122–129. https://doi.org/10.1016/j.anireprosci.2011.04.009

Beirão, J., Soares, F., Herráez, M.P., Dinis, M.T., Cabrita, E., 2009. Sperm quality evaluation in Solea senegalensis during the reproductive season at cellular level. Theriogenology 72, 1251–1261. https://doi.org/10.1016/j.theriogenology.2009.07.021

Bombardelli, R.A., Sanches, E.A., Baggio, D.M., Sykora, R.M., Souza, B.E. de, Tessaro, L., Piana, P.A., 2013. Effects of the spermatozoa: oocyte ratio, water volume and water temperature on artificial fertilization and sperm activation of cascudo-preto. Rev. Bras. Zootec. 42, 1–6. https://doi.org/10.1590/S1516-35982013000100001

Bromage, N., 1992. Propagation and stock improvement, in: Bromage, N., Shepherd, J. (Eds.), Intensive Fish Farming. Blackwell Science, Oxford, UK, pp. 103–153.

Brown, N., 2010. Halibut aquaculture in North America, in: Daniels, H., Watanabe, W. (Eds.), Practical FLatfish Culture and Stock Enhancement. Blackwell Publishing, Ames, Iowa, USA.

Butts, I., Roustaian, P., Litvak, M., 2012. Fertilization strategies for winter flounder: effects of spermatozoa density and the duration of gamete receptivity. Aquat. Biol. 16, 115–124. https://doi.org/10.3354/ab00439

Butts, I.A.E., Trippel, E.A., Litvak, M.K., 2009. The effect of sperm to egg ratio and gamete contact time on fertilization success in Atlantic cod Gadus morhua L. Aquaculture 286, 89–94. https://doi.org/10.1016/j.aquaculture.2008.09.005

Cabrita, E., Soares, F., Beirão, J., García-López, A., Martínez-Rodríguez, G., Dinis, M.T., 2011. Endocrine and milt response of Senegalese sole, Solea senegalensis, males maintained in captivity. Theriogenology 75, 1–9. https://doi.org/10.1016/j.theriogenology.2010.07.003

Cabrita, E., Soares, F., Dinis, M.T., 2006. Characterization of Senegalese sole, Solea senegalensis, male broodstock in terms of sperm production and quality. Aquaculture 261, 967–975. https://doi.org/10.1016/j.aquaculture.2006.08.020

Carazo, I., Chereguini, O., Martín, I., Huntingford, F., Duncan, N., 2016. Reproductive ethogram and mate selection in captive wild Senegalese sole (Solea senegalensis). Span. J. Agric. Res. 14, e0401. https://doi.org/10.5424/sjar/2016144-9108

Casselman, S.J., Schulte-Hostedde, A.I., Montgomerie, R., 2006. Sperm quality influences male fertilization success in walleye (*Sander vitreus*). Can. J. Fish. Aquat. Sci. 63, 2119–2125. https://doi.org/10.1139/f06-108

Cejko, B.I., Żarski, D., Targońska, K., Krejszeff, S., Kucharczyk, D., Glogowski, J., 2010. Osmolality of Seminal Plasma as an Indicator of Milt Contamination with Urine Based on the Example of the Tench *Tinca tinca* (L.). Pol. J. Nat. Sci. 25, 287–298. https://doi.org/10.2478/v10020-010-0026-6

Chauvigné, F., González, W., Ramos, S., Ducat, C., Duncan, N., Giménez, I., Cerdà, J., 2018. Seasonal-and dose-dependent effects of recombinant gonadotropins on sperm production and quality in the flatfish Solea senegalensis. Comp. Biochem. Physiol. A. Mol. Integr. Physiol. 225, 59–64. https://doi.org/10.1016/j.cbpa.2018.06.022

Chauvigné, F., Ollé, J., González, W., Duncan, N., Giménez, I., Cerdà, J., 2017. Toward developing recombinant gonadotropin-based hormone therapies for increasing fertility in the flatfish Senegalese sole. PLOS ONE 12, e0174387. https://doi.org/10.1371/journal.pone.0174387

Chereguini, O., de la Banda, I.G., Rasines, I., Fernandez, A., 1999. Artificial fertilization in turbot, Scophthalmus maximus (L.): different methods and determination of the optimal sperm-egg ratio. Aquac. Res. 30, 319–324. https://doi.org/10.1046/j.1365-2109.1999.00326.x

Domeier, M.L., Colin, P.L., 1997. Tropical reef fish spawning aggregations: defined and reviewed. Bull. Mar. Sci. 60, 698–726.

Duncan, N., Carazo, I., Chereguini, O., Mañanós, E., 2019. Mating Behaviour, in: Munoz-Cueto, J., Mañanós-Sánchez, E., Sánchez-Vázquez, J. (Eds.), Biology of Sole. CRC Press, Boca Raton.

Fatsini, E., Carazo, I., Chauvigné, F., Manchado, M., Cerdà, J., Hubbard, P.C., Duncan, N.J., 2017. Olfactory sensitivity of the marine flatfish *Solea senegalensis* to conspecific body fluids. J. Exp. Biol. 220, 2057–2065. https://doi.org/10.1242/jeb.150318

Fatsini, E., González, W., Ibarra-Zatarain, Z., Napuchi, J., Duncan, N.J., 2020. The presence of wild Senegalese sole breeders improves courtship and reproductive success in cultured conspecifics. Aquaculture 519, 734922. https://doi.org/10.1016/j.aquaculture.2020.734922

Gallego, V., Carneiro, P.C.F., Mazzeo, I., Vílchez, M.C., Peñaranda, D.S., Soler, C., Pérez, L., Asturiano, J.F., 2013a. Standardization of European eel (Anguilla anguilla) sperm motility evaluation by CASA software. Theriogenology 79, 1034–1040. https://doi.org/10.1016/j.theriogenology.2013.01.019

Gallego, V., Pérez, L., Asturiano, J.F., Yoshida, M., 2013b. Relationship between spermatozoa motility parameters, sperm/egg ratio, and fertilization and hatching rates in pufferfish (Takifugu niphobles). Aquaculture 416–417, 238–243. https://doi.org/10.1016/j.aquaculture.2013.08.035

García-López, Á., Martínez-Rodríguez, G., Sarasquete, C., 2005. Male reproductive system in Senegalese sole Solea senegalensis (Kaup): Anatomy, histology and histochemistry. Histol. Histopathol. 20, 1179–1189. https://doi.org/10.14670/HH-20.1179

Gibson, R.N., Stoner, A.W., Ryer, C.H., 2014. The behaviour of flatfishes, in: Gibson, R.N., Nash, R.D.M., Geffen, A.J., van der Veer, H.W. (Eds.), Flatfishes: Biology and Exploitation.

González-López, W.Á., Ramos-Júdez, S., Giménez, I., Duncan, N.J., 2020. Sperm contamination by urine in Senegalese sole (Solea senegalensis) and the use of extender solutions for short-term chilled storage. Aquaculture 516, 734649. https://doi.org/10.1016/j.aquaculture.2019.734649

Guzmán, J.M., Cal, R., García-López, Á., Chereguini, O., Kight, K., Olmedo, M., Sarasquete, C., Mylonas, C.C., Peleteiro, J.B., Zohar, Y., Mañanós, E.L., 2011. Effects of in vivo treatment with the dopamine antagonist pimozide and gonadotropin-releasing hormone agonist (GnRHa) on the reproductive axis of Senegalese sole (Solea senegalensis). Comp. Biochem. Physiol. A. Mol. Integr. Physiol. 158, 235–245. https://doi.org/10.1016/j.cbpa.2010.11.016

Guzmán, J.M., Norberg, B., Ramos, J., Mylonas, C.C., Mañanós, E.L., 2008. Vitellogenin, steroid plasma levels and spawning performance of cultured female Senegalese sole (Solea senegalensis). Gen. Comp. Endocrinol. 156, 285–297. https://doi.org/10.1016/j.ygcen.2008.02.002

Guzmán, J.M., Ramos, J., Mylonas, C.C., Mañanós, E.L., 2009. Spawning performance and plasma levels of GnRHa and sex steroids in cultured female Senegalese sole (Solea senegalensis) treated with different GnRHa-delivery systems. Aquaculture 291, 200–209. https://doi.org/10.1016/j.aquaculture.2009.03.024

Ibarra-Zatarain, Z., Duncan, N., 2015. Mating behaviour and gamete release in gilthead seabream (Sparus aurata, Linnaeus 1758) held in captivity. Span. J. Agric. Res. 13, e0401. https://doi.org/10.5424/sjar/2015131-6750

Kholodnyy, V., Gadêlha, H., Cosson, J., Boryshpolets, S., 2020. How do freshwater fish sperm find the egg? The physicochemical factors guiding the gamete encounters of externally fertilizing freshwater fish. Rev. Aquac. 12, 1165–1192. https://doi.org/10.1111/raq.12378

Linhart, O., Rodina, M., Bastl, J., Cosson, J., 2003. Urinary bladder, ionic composition of seminal fluid and urine with characterization of sperm motility in tench (Tinca tinca L.). J. Appl. Ichthyol. 19, 177–181. https://doi.org/10.1046/j.1439-0426.2003.00470.x

Liu, Xin-fu, Liu, Xue-zhou, Lian, J., Wang, Y., Zhang, F., Yu, H., Ma, A., Liu, S., Zhai, J., 2008. Large scale artificial reproduction and rearing of Senegal sole, Solea senegalensis Kaup. Mar. Fish. Res. 29, 10–16.

Mañanós, E., Duncan, N., Mylonas, C., 2008. Reproduction and Control of Ovulation, Spermiation and Spawning in Cultured Fish, in: Cabrita, E., Robles, V., Herráez, P. (Eds.), Methods in Reproductive Aquaculture, Marine Biology. CRC Press, pp. 3–80. https://doi.org/10.1201/9780849380549.sec1

Marrero-Alemán, C., González-López, W., Ramos-Júdez, S., Navarro, I., Duncan, N., 2019. Artificial fertilisation in Senegalese sole (Solea senegalensis): induction with GnRHa and determination of egg quality., in: Aquaculture Europe 19, Abstracts. Presented at the Aquaculture Europe 19, European Aquaculture Society, Berlin, Germany.

Martín, I., Carazo, I., Rasines, I., Rodríguez, C., Fernández, R., Martínez, P., Norambuena, F., Chereguini, O., Duncan, N., 2020. Reproductive performance of captive Senegalese sole, Solea senegalensis, according to the origin (wild or cultured) and gender. Span. J. Agric. Res. 17, e0608. https://doi.org/10.5424/sjar/2019174-14953

Martín, I., Rasines, I., Gómez, M., Rodríguez, C., Martínez, P., Chereguini, O., 2014. Evolution of egg production and parental contribution in Senegalese sole, Solea senegalensis, during four consecutive spawning seasons. Aquaculture 424–425, 45–52. https://doi.org/10.1016/j.aquaculture.2013.12.042

Moccia, R.D., Munkittrick, K.R., 1987. Relationship between the fertilization of rainbow trout (Salmo gairdneri) eggs and the motility of spermatozoa. Theriogenology 27, 679–688. https://doi.org/10.1016/0093-691X(87)90061-6

Morais, S., Aragão, C., Cabrita, E., Conceição, L.E.C., Constenla, M., Costas, B., Dias, J., Duncan, N., Engrola, S., Estevez, A., Gisbert, E., Mañanós, E., Valente, L.M.P., Yúfera, M., Dinis, M.T., 2016. New developments and biological insights into the farming of *Solea senegalensis* reinforcing its aquaculture potential. Rev. Aquac. 8, 227–263. https://doi.org/10.1111/raq.12091

Mylonas, C.C., Duncan, N.J., Asturiano, J.F., 2017. Hormonal manipulations for the enhancement of sperm production in cultured fish and evaluation of sperm quality. Aquaculture 472, 21–44. https://doi.org/10.1016/j.aquaculture.2016.04.021

Norambuena, F., Estevez, A., Bell, G., Carazo, I., Duncan, N., 2012a. Proximate and fatty acid compositions in muscle, liver and gonads of wild versus cultured broodstock of Senegalese sole (Solea senegalensis). Aquaculture 356–357, 176–185. https://doi.org/10.1016/j.aquaculture.2012.05.018

Norambuena, F., Estévez, A., Mañanós, E., Bell, J.G., Carazo, I., Duncan, N., 2013a. Effects of graded levels of arachidonic acid on the reproductive physiology of Senegalese sole (Solea senegalensis): Fatty acid composition, prostaglandins and steroid levels in the blood of broodstock bred in captivity. Gen. Comp. Endocrinol. 191, 92–101. https://doi.org/10.1016/j.ygcen.2013.06.006

Norambuena, F., Estévez, A., Sánchez-Vázquez, F.J., Carazo, I., Duncan, N., 2012b. Self-selection of diets with different contents of arachidonic acid by Senegalese sole (Solea senegalensis) broodstock. Aquaculture 364–365, 198–205. https://doi.org/10.1016/j.aquaculture.2012.08.016

Norambuena, F., Mackenzie, S., Bell, J.G., Callol, A., Estévez, A., Duncan, N., 2012c. Prostaglandin (F and E, 2- and 3-series) production and cyclooxygenase (COX-2) gene expression of wild and cultured broodstock of senegalese sole (Solea senegalensis). Gen. Comp. Endocrinol. 177, 256–262. https://doi.org/10.1016/j.ygcen.2012.04.009

Norambuena, F., Morais, S., Estévez, A., Bell, J.G., Tocher, D.R., Navarro, J.C., Cerdà, J., Duncan, N., 2013b. Dietary modulation of arachidonic acid metabolism in senegalese sole (Solea Senegalensis) broodstock reared in captivity. Aquaculture 372–375, 80–88. https://doi.org/10.1016/j.aquaculture.2012.10.035

Parker, G.A., Pizzari, T., 2010. Sperm competition and ejaculate economics. Biol. Rev. 85, 897–934. https://doi.org/10.1111/j.1469-185X.2010.00140.x

Perchec Poupard, G., Paxion, C., Cosson, J., Jeulin, C., Fierville, F., Billard, R., 1998. Initiation of carp spermatozoa motility and early ATP reduction after milt contamination by urine. Aquaculture 160, 317–328. https://doi.org/10.1016/S0044-8486(97)00301-3

Prins, J., 2012. Product and Process Comparions, in: Croarkin, C., Tobias, P. (Eds.), NIST/SEMATECH e-Engineering Statistics Handbook. http://www.itl.nist.gov/div898/handbook/ https://doi.org/10.18434/M32189 Accessed 06/08/2020

Ramos-Júdez, S., González, W., Dutto, G., Mylonas, C.C., Fauvel, C., Duncan, N., 2019. Gamete quality and management for in vitro fertilisation in meagre (Argyrosomus regius). Aquaculture 509, 227–235. https://doi.org/10.1016/j.aquaculture.2019.05.033

Rasines, I., Gómez, M., Martín, I., Rodríguez, C., Mañanós, E., Chereguini, O., 2013. Artificial fertilisation of cultured Senegalese sole (Solea senegalensis): Effects of the time of day of hormonal treatment on inducing ovulation. Aquaculture 392–395, 94–97. https://doi.org/10.1016/j.aquaculture.2013.02.011

Rasines, I., Gómez, M., Martín, I., Rodríguez, C., Mañanós, E., Chereguini, O., 2012. Artificial fertilization of Senegalese sole (Solea senegalensis): Hormone therapy administration methods, timing of ovulation and viability of eggs retained in the ovarian cavity. Aquaculture 326–329, 129–135. https://doi.org/10.1016/j.aquaculture.2011.11.021

Riesco, M.F., Valcarce, D.G., Martínez-Vázquez, J.M., Martín, I., Calderón-García, A.Á., Gonzalez-Nunez, V., Robles, V., 2019. Male reproductive dysfunction in Solea senegalensis: new insights into an unsolved question. Reprod. Fertil. Dev. 31, 1104. https://doi.org/10.1071/RD18453

Rizzo, E., Godinho, H.P., Sato, Y., 2003. Short-term storage of oocytes from the neotropical teleost fish Prochilodus marggravii. Theriogenology 60, 1059–1070. https://doi.org/10.1016/S0093-691X(03)00108-0

Samarin, A.M., Amiri, B.M., Soltani, M., Mohammad, R., Kamali, A., Naghavi, M.R., 2011. Effects of storage duration and storage temperature on viability of stored ova of kutum (Rutilus frisii kutum) in ovarian fluid. Afr. J. Biotechnol. 10, 12309–12314. https://doi.org/10.5897/AJB11.919

Sanches, E.A., Caneppele, D., Okawara, R.Y., Damasceno, D.Z., Bombardelli, R.A., Romagosa, E., 2016. Inseminating dose and water volume applied to the artificial fertilization of Steindachneridion parahybae (Steindachner, 1877) (Siluriformes: Pimelodidae): Brazilian endangered fish. Neotropical Ichthyol. 14, e140158. https://doi.org/10.1590/1982-0224-20140158

Sarosiek, B., Dryl, K., Krejszeff, S., Żarski, D., 2016. Characterization of pikeperch (Sander lucioperca) milt collected with a syringe and a catheter. Aquaculture 450, 14–16. https://doi.org/10.1016/j.aquaculture.2015.06.040

Stockley, P., Gage, M.J.G., Parker, G.A., Møller, A.P., 1997. Sperm Competition in Fishes: The Evolution of Testis Size and Ejaculate Characteristics. Am. Nat. 149, 933–954. https://doi.org/10.1086/286031

Suquet, M., Billard, R., Cosson, J., Normant, Y., Fauvel, C., 1995. Artificial insemination in turbot (Scophthalmus maximus): determination of the optimal sperm to egg ratio and time of gamete contact. Aquaculture 133, 83–90. https://doi.org/10.1016/0044-8486(94)00395-5

Tatarenkov, A., Barreto, F., Winkelman, D.L., Avise, J.C., 2006. Genetic Monogamy in the Channel Catfish, Ictalurus Punctatus, a Species with Uniparental Nest Guarding. Copeia 2006, 735–741. https://doi.org/10.1643/0045-8511(2006)6[735:GMITCC]2.0.CO;2

Yanagimachi, R., Harumi, T., Matsubara, H., Yan, W., Yuan, S., Hirohashi, N., Iida, T., Yamaha, E., Arai, K., Matsubara, T., Andoh, T., Vines, C., Cherr, G.N., 2017. Chemical and physical guidance of fish spermatozoa into the egg through the micropyle. Biol. Reprod. 96, 780–799. https://doi.org/10.1093/biolre/iox015

